# Spreading depolarization causes reversible neuronal mitochondria fragmentation and swelling in healthy, normally perfused neocortex

**DOI:** 10.1101/2024.01.22.576364

**Authors:** Jeremy Sword, Ioulia V. Fomitcheva, Sergei A. Kirov

## Abstract

Mitochondrial function is tightly linked to their morphology, and fragmentation of dendritic mitochondria during noxious conditions suggests loss of function. In the normoxic cortex, spreading depolarization (SD) is a phenomenon underlying migraine aura. It is unknown whether mitochondria structure is affected by normoxic SD. *In vivo* two-photon imaging followed by quantitative serial section electron microscopy (ssEM) was used to monitor dendritic mitochondria in the normoxic cortex of urethane-anesthetized mature male and female mice during and after SD initiated by focal KCl microinjection. Structural dynamics of dendrites and their mitochondria were visualized by transfecting excitatory, glutamatergic neurons of the somatosensory cortex with bicistronic AAV, which induced tdTomoto labeling in neuronal cytoplasm and mitochondria labeling with roGFP. Normoxic SD triggered a rapid fragmentation of dendritic mitochondria alongside dendritic beading, both reversible; however, mitochondria took significantly longer to recover. Several rounds of SD resulted in transient mitochondrial fragmentation and dendritic beading without accumulating injury, as both recovered. SsEM corroborated normoxic SD-elicited dendritic and mitochondrial swelling and transformation of the filamentous mitochondrial network into shorter, swollen tubular and globular structures. Our results revealed normoxic SD-induced disruption of the dendritic mitochondrial structure that might impact mitochondrial bioenergetics during migraine with aura.

## Introduction

Comprehensive clinical data reveal that spreading depolarization (SD) is central in developing cortical lesions in patients with acute brain injury^1–5^. SD waves course across the cortex at a velocity of 2-9 mm/min^6, 7^, depolarizing neurons and glia and eliciting the collapse of transmembrane ion gradients, leading to cytotoxic edema and distortion of synaptic circuitry^8–10^. Common biophysical features characterize a continuous spectrum of SD waves ranging from long-lasting detrimental events under severe ischemia to short-lasting harmless events in the healthy brain^2, 11, 12^. Thus, on the one end of the spectrum, SD is thought to facilitate neuronal death in the metabolically compromised brain^13–15^, but on the opposite end in the non-injured brain, SD, which is assumed to constitute the neurological basis of migraine with aura^8, 16, 17^ is innocuous^18^.

Still, regardless of the energy state of the cortical tissue invaded by SD, cytotoxic edema rapidly develops^19–22^ following intracellular water influx as a consequence of SD-induced massive net gain of excess electrolytes in the cytoplasm^12, 23^. Focal dendritic swelling (also referred to as dendritic beading) is easily detectable in real-time by two-photon laser scanning microscopy (2PLSM) and, therefore, serves as a reliable read-out of the onset of neuronal cytotoxic edema^24–26^. Dendrites swell along the entire spectrum of SD waves. Short-lasting SD elicits transient beading in healthy, normally perfused cortex^24, 27–29^. Dendritic beading is also reversible in the ischemic penumbra during intermediate SD, but clusters of SDs facilitate terminal dendritic beading, contributing to the ischemic core expansion into the penumbra^30, 31^. Likewise, long-lasting SD during severe global ischemia and without immediate reperfusion results in terminal beading^2, 25^. Consequently, cytotoxic edema is widely accepted as an essential contributor to acute, irreversible brain injury^32, 33^.

Long-term neuronal survival in ischemic disorders is tightly linked with mitochondrial function^34^. Mitochondrial fragmentation under pathological conditions was shown to be accompanied by mitochondrial oxidation and reduced mitochondrial membrane potential^35^, thus providing “proxy” functional read-outs for diminished bioenergetics of morphologically altered mitochondria *in vivo*. Dendrites of principal neurons contain long filamentous mitochondria^22, 36, 37^. Using real-time *in vivo* two-photon laser scanning microscopy (2PLSM), we have shown that mitochondrial structure is very sensitive to the noxious conditions leading to brain damage^37^. A rapid fragmentation of neocortical dendritic mitochondria during mild ischemic and traumatic injury was reversible, highlighting mitochondria as a potential therapeutic target.

Likewise, our recent *in vivo* 2PLSM and ultrastructural study revealed reversible fragmentation of dendritic mitochondria alongside transient dendritic swelling elicited by short-lasting SD during transient ischemic attack^22^. SD is the pathophysiological correlate of the migraine aura, and SD in the normoxic cortex simulates this condition. The dynamics of *in vivo* mitochondrial response to normoxic SD is unknown. It is unknown if normoxic SD fragments dendritic mitochondria and, if so, how fast mitochondrial fusion occurs during repolarization. Based on the framework of the stroke-migraine depolarization continuum^2, 11, 12^, we hypothesized that, like ischemia-induced SD, normoxic SD triggers rapid and readily reversible mitochondrial swelling and fragmentation in the healthy, normally perfused cortex. Here, *in vivo* 2PLSM followed by quantitative serial section electron microscopy (ssEM) was used to assess the integrity of mitochondria and dendrites in the somatosensory cortex of urethane-anesthetized mice during normoxic SD evoked by focal KCl microinjection.

## Materials and Methods

### Animals and surgical procedures

All procedures followed National Institutes of Health guidelines for the humane care and use of laboratory animals and underwent yearly review by the Animal Care and Use Committee at the Medical College of Georgia. All experiments were performed in accordance with ARRIVE guidelines^38^. The mice were bred and housed in group cages in the certified animal facilities in a 12 h light/dark cycle, with constant temperature (22±1°C), and provided with food and water ad libitum. A total of 39 C57BL/6J mice (JAX stock #000664) of both sexes at an average age of 6 months were used. For viral transduction, mice at postnatal day 21 were anesthetized with isoflurane (4% induction, 1–1.5% maintenance in 21% or 100% oxygen), and a small hole was made over the somatosensory cortex centered at −1.8 mm from bregma and 2.8 mm lateral to allow adeno-associated virus (AAV) delivery. Mice were injected with 250 nL AAV/DJ-CamKII(0.4)-mito-roGFP1-T2A-tdTomato-WPRE; titer 4.5 x 10^12^ GC/mL, undiluted (Vector Biolabs, custom vector). AAV expression labeled the cytoplasm with tdTomato and mitochondria with roGFP1 fused to the mitochondrial-targeting sequence of human pyruvate dehydrogenase alpha 1^39^. Viruses were injected at 100 nL/min using micropump (WPI) with micropipettes (Drummond) placed to a depth of ∼500 µm, and mice were used at least two months later.

For intravital imaging, mice were anesthetized with an intraperitoneal injection of urethane (1.5 mg/g), the trachea was cannulated, and animals were ventilated with SAR-1000 (CWE). End-tidal CO_2_ was monitored with Capnoscan (Kent Scientific) using relative values previously validated with blood gas monitoring in a separate cohort of animals to maintain etCO_2_ in a normal physiological range of 37±2 mmHg. Body temperature was maintained at 37±1°C. The depth of anesthesia, blood oxygen saturation level (>90%), and heart rate (450–650 beats/min) were monitored with a MouseOx pulse oximeter (STARR Life Sciences). Hydration was maintained by intraperitoneal injection of 200 µL 0.9% NaCl with 20 mM glucose. Implantation of the cranial window followed standard protocol^30^. An optical chamber ∼4 mm in diameter centered at stereotaxic coordinates −1.8 mm from bregma and 2.8 mm lateral over the somatosensory cortex was constructed by covering the intact dura with a thin layer of 1.5% agarose prepared in a cortex buffer (in mM: 135 NaCl, 5.4 KCl, 1 MgCl_2_, 1.8 CaCl_2_, and 10 HEPES, pH 7.3). The chamber was open for access with glass micropipettes. The Ag/AgCl pellet ground electrode (A-M Systems) was sutured under the skin above the nasal bone. All chemicals were from Sigma unless indicated otherwise.

### Electrophysiology and SD triggering

The DC potential and spontaneous electrocorticographic (ECoG) activity were recorded with a glass microelectrode (1–2 MΩ) inserted next to imaged dendrites within layer I of the somatosensory cortex. SDs were evoked by focal pressure injection of <5 nl of 1 M KCl with a Picospritzer (Parker Hannifin) using a glass micropipette inserted to the depth of 200–300 μm opposite the recording electrode and outside of the imaging area, preventing direct KCl administration. Signals were recorded with a MultiClamp 700B amplifier, filtered at 1 kHz, digitized at 10 kHz with Digidata 1550, and analyzed with pClamp 10 (Molecular Devices).

### Imaging modalities, processing, and analysis

The Nikon A1R MP multiphoton system mounted on the FN1 upright microscope was used for imaging with a 25x/1.1 NA water-immersion objective. The Spectra-Physics Mai-Tai eHP-FS DeepSee laser tuned to 910 nm was used for two-photon excitation, and emission light was measured by GaAsP detectors using bandpass filters (500–550 nm) and (575-625 nm). Image z-stacks were acquired from 60 µm to the surface at 1 μm increments across a 75×75 μm imaging field (512×512 pixels).

The Fiji image processing package distribution of ImageJ (NIH) was used for image analysis and processing. Huygens Professional image deconvolution software (Scientific Volume Imaging) was used to process images in figures. Focal swelling (beading) of dendrites, which resembles beads on a string, was identified by rounded regions extending beyond the initial diameter of the dendrite. The progression of dendritic beading (swelling) and fragmentation of the mitochondrial network was assessed manually by quantifying the amount of dendritic beading or mitochondrial fragmentation in an imaging field following published protocols^25, 28, 40^. Because the resolution and signal-to-noise ratio always present a challenge for analyses of small structures such as mitochondria, images were not randomized to keep track of individual mitochondria in the image sequences. Briefly, individual image planes were randomly selected from the middle of a stack, matched with corresponding planes before, during, and immediately after SD, and divided into 12×12 squares (6×6 µm). Only the squares containing dendrites or mitochondria were counted, and the percentage of squares containing beaded dendrites or fragmented mitochondria was calculated. Maximum diameters of mitochondria and dendrites were assessed from random segments of 10 mitochondria and dendrites in each imaging field immediately before, during, and after SD to further estimate the speed of morphological changes and recovery.

### Electron microscopy and analysis

Mice were perfusion-fixed through the heart at 180 mmHg with mixed aldehydes (2% paraformaldehyde, 2.5% glutaraldehyde, 2 mM CaCl_2_, 4 mM MgSO_4_, in 0.1 M sodium cacodylate, pH 7.4). The area of intravital imaging was identified in the fixed brain with 2PLSM using the blood vasculature and the dendritic pattern as a guide.

Then, the whole fixed brain was removed and embedded in an agar block, and the imaging area was labeled with 100 µm laser marks (“brands”) on the cortical surface using a Ti:sapphire laser beam^41, 42^. Coronal slices 100 µm thick were cut, and sections with laser marks were identified. Additional laser brands were added to the fixed coronal slices with the intravital imaging area halfway between laser brands. Tissue was processed with standard microwave-enhanced procedures through osmium, uranyl acetate, dehydration with a graded ethanol series, and embedding in Epon–Araldite resin standard^43^. Using the laser brands as a guide, two series containing 136 and 194 sections, each ∼53 nm thick, were cut with a diamond knife on an EM UC6 ultramicrotome (Leica) at a depth of intravital imaging from two mice perfusion-fixed within 2 min of SD onset. Three series containing 190, 203, and 204 sections, each ∼55 nm thick, were cut at a depth of intravital imaging from two *Sham* mice. Sections were collected on pioloform-coated copper Synaptek slot grids (Electron Microscopy Sciences) and stained with uranyl acetate and lead citrate. All chemicals for EM were from Electron Microscopy Sciences except CaCl_2_ and MgSO_4_, which were from Fisher

Scientific. Serial sections were photographed at 5000x with the JEOL 1230 transmission electron microscope using the UltraScan 4000 camera (Gatan). The 3D alignment, surfaced reconstructions, and analyses, blind as to condition, were completed using RECONSTRUCT software^44^. Pixel size was calibrated with a diffraction grating replica (Ernest F. Fullam). Section thickness was calculated by dividing the diameters of longitudinally sectioned mitochondria by the number of sections they occupied^45^.

### Tissue pO_2_ Measurements

A Clark-type polarographic oxygen microelectrode (OX-10; <10 μM tip; Unisense) with a guard cathode was used for tpO_2_ measurements, with a sensitivity sphere twice the tip diameter. Each electrode was calibrated before experiments in air-saturated 0.9% saline and zero oxygen solution of 0.1 M sodium L-ascorbate plus 0.1 M NaOH in saline. The oxygen electrode was inserted into the somatosensory cortex at the same depth as the microelectrode for ECoG recordings and 20-60 µm away. The oxygen electrodes were connected to a high-impedance picoammeter (Oxy-meter; Unisense) that measured the electrode currents, A/D converted, and recorded at 1 or 50 Hz for display in SensorTrace (Unisense).

### Statistical analyses

Microsoft Excel was used to organize and plot data. SigmaStat (Systat) and GraphPad Prism were used for statistical analyses. Normality was tested with the Shapiro–Wilk test. Parametric data were analyzed with two-tailed paired Student’s t-test and one-way repeated-measures analysis of variance (RM ANOVA) followed by Tukey’s post hoc test. Data significantly deviant from normal distribution were analyzed using the Kolmogorov–Smirnov (K–S) test, Wilcoxon signed rank test, Kruskal–Wallis ANOVA on ranks followed by Dunn’s post hoc test and Friedman RM ANOVA on ranks followed by Dunn’s or Student-Newman-Keuls (SNK) post hoc tests. Group sizes were determined by statistical power analyses based on estimates from preliminary (or published) studies and aiming to detect 20% change with the significance level set at α=0.05, study power set at 80%, and standard deviation (SD) based on experimental variability. The linear regression analysis and the Pearson correlation coefficient were calculated to quantify the strength of the relationship between the two variables. The slopes of the regression lines were compared using the homogeneity-of-slopes model. A χ2 test was used to analyze data arranged in contingency tables. The sample size of each experimental group is given in the “Results” and “Figure legends.” Parametric data are presented as mean ± SD. Nonparametric data are presented as the median and interquartile range (IQR), i.e., the difference between the third and the first quartile. The significance criterion was set at *P*<0.05.

## Results

### Rapid fragmentation of dendritic mitochondria alongside dendritic beading during normoxic SD *in vivo*

We used AAV carrying mito-roGFP-T2A-tdTomato construct under the control of the CamKII promoter to visualize mitochondria and dendrites of excitatory, glutamatergic neurons in the somatosensory cortex of mice with *in vivo* 2PLSM. Pressure injection of <5 nl of 1M KCl in healthy, normally perfused cortex away from the site of imaging evoked SD, which invaded the imaging field as was detected by the large negative shift of the cortical slow DC potential (**Fig. 1A**). As expected, normoxic SD induced rapid dendritic beading^24, 27–29^ revealing the onset of neuronal cytotoxic edema^21, 22^. Remarkably, besides dendritic beading, normoxic SD triggered swift mitochondrial fragmentation (**Fig. 1B**). Indeed, the fraction of fragmented mitochondria in the imaging field has significantly increased during SD propagation in 33 out of 36 experimental animals (χ^2^-test for each mouse). In comparison, significant dendritic beading during SD was detected in 35 mice (χ^2^-tests). As summarized in **Figure 1C** for all 36 experimental animals, the percent of mitochondrial fragmentation in the imaging field significantly increased from 12.0% (IQR: 5.1-19.5%) at baseline to 98.0% (IQR: 89.7-100%) during SD (P<0.001, Wilcoxon signed rank test). In parallel, the percent of dendritic beading significantly increased from 6.6% (IQR: 2.6-9.6%) at baseline to 99.6% (IQR: 92-100%) during SD (P<0.001, Wilcoxon signed rank test).

**Figure 1.**
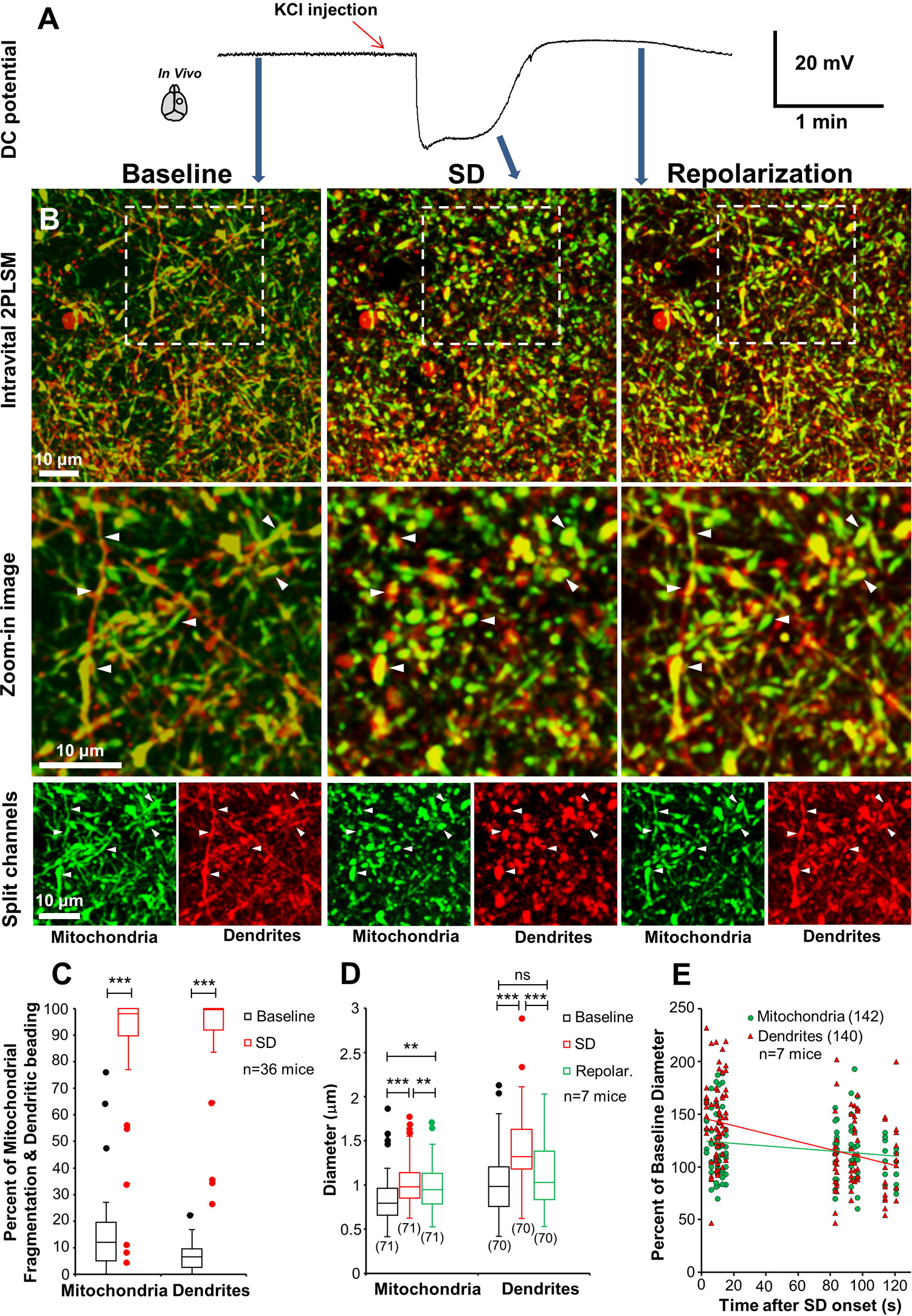
*In vivo* dendritic mitochondria dynamics during normoxic SD. **A,** Recording of the DC cortical potential from a glass microelectrode in the location of 2PLSM imaging in the layer I of the somatosensory cortex of a mouse injected with AAV-CamKII-mito-roGFP-T2A-tdTomato vector. The large negative deflection in the DC potential indicates SD triggered by KCl injection (red arrow) away from the imaging site. Blue arrows mark the exact time points on the SD recording when intravital 2PLSM images in (B) were acquired. **B,** Large-scale dual-color images of mitochondria (green) and dendrites (red) shown at the top row and corresponding zoom-in images of the white boxed areas in the middle row illustrate mitochondrial fragmentation and dendritic beading during normoxic SD. The dual-color images are split at the bottom row to show mitochondria and dendrites in separate channels. Arrowheads mark the same spots on mitochondria and dendrites in the image sequences. **C**, Quantification of SD-induced mitochondrial fragmentation and dendritic beading in the imaging field immediately after SD onset (***P<0.001, Wilcoxon Signed Rank Tests). Circles depict outliers. **D,** Box and Whisker plot of changes in the diameter of mitochondria and dendrites during the onset of SD and repolarization. Mitochondria significantly swelled at the SD onset and were still swollen during or immediately after repolarization (χ2_(2)_=43.19, P<0.0001, Friedman RM ANOVA on Ranks with Dunn’s post hoc test. ***P<0.0001, **P<0.01). Dendrites were beaded/swollen by SD but completely recovered during repolarization (χ2_(2)_=64.31, P<0.0001, Friedman RM ANOVA on Ranks with Dunn’s post hoc test. ***P<0.0001). Circles depict outliers. Numbers in parentheses indicate the number of mitochondria and dendrites included in the calculations. **E,** Rate of the recovery from SD-induced swelling measured at the SD onset and during repolarization was faster for dendrites than for mitochondria, as indicated by the significant difference in the slopes of regression lines (F_(1, 278)_=13.71, P<0.001, the homogeneity-of-slopes model for two independent samples).

Next, in 7 mice, we assessed SD-induced swelling and recovery of mitochondria and dendrites by measuring the diameter of randomly selected dendrites and their mitochondria organelles. All SDs were comparable between animals in amplitude (18.4 ± 2.8 mV) and duration (93 ± 20.7 s, as measured at half maximum of the DC amplitude) and triggered significant mitochondrial and dendritic swelling (**Fig. 1D**; P<0.0001 for both structures, Friedman RM ANOVA on Ranks). At 9.6 ± 4.1 s after SD onset, the mitochondrial diameter increased by 18.5% (IQR: 0-44.4%) (P<0.0001, Dunn’s post hoc test, relative to the baseline). At the same time, the dendritic diameter was larger by 40.8% (IQR: 20.2-66.1%) (P<0.0001, Dunn’s post hoc test). At 98.1 ± 14.4 s after SD onset, i.e., during repolarization or immediately after, the dendritic diameter swiftly recovered to 3.8% (IQR: −14.9-31.0%) of baseline values (P=0.3, Dunn’s post hoc test), while mitochondria were still significantly swollen by 13.0% (IQR: −7.1-28.3%) (P<0.01, Dunn’s post hoc test). Indeed, the linear regression analyses (**Fig. 1E**) revealed a moderate negative correlation between the change in dendritic diameter immediately after SD onset and during repolarization (P<0.0001; R=-0.43, Pearson correlation), signifying a fast decrease in the size of dendritic beads. In contrast, the negative correlation between the reduction in mitochondrial diameter and the time after SD onset was very weak, albeit significant (P=0.03; R=-0.18, Pearson correlation), demonstrating slower recovery. Accordingly, the comparison of the slopes of regression lines indicated that dendrites recovered faster (**Fig. 1E**; P<0.001, the homogeneity-of-slopes model for two independent samples). These results demonstrate that normoxic SD triggered dendritic mitochondria fragmentation alongside dendritic beading. Dendrites recovered during repolarization, but mitochondrial recovery was slower.

### SD-induced dendritic mitochondria fragmentation during several rounds of normoxic SD

Several rounds of normoxic SD *in vivo* do not result in accumulating dendritic beading and injury as dendrites swiftly recover between SDs^27, 29^. Under noxious conditions, fragmentation of the mitochondrial network into small spherical structures is considered a hallmark of mitochondrial injury^37, 46^. Consequently, since mitochondrial recovery from normoxic SD-induced fragmentation was slower than dendritic recovery, we tested whether mitochondrial fusion would be complete between subsequent normoxic SDs and whether several SDs would result in irreversible fragmentation.

Three SDs elicited in 5 mice at 23 min (IQR: 20-33 min) apart (**Fig. 2A** bottom panels for representative SD traces) were comparable in duration (**Fig. 2B**; P=0.26, RM ANOVA) but different in amplitude (**Fig. 2C**; P<0.005, RM ANOVA). Each SD triggered transient mitochondrial fragmentation that recovered by the time when the next SD was elicited (**Fig. 2A**). Indeed, quantification revealed that the fraction of fragmented mitochondria in the imaging field increased from 6.1 ± 2.8% at baseline to 98.2±1.9%, 99±1.7% and 95.8±4.9% during first, second and third SD respectfully (F_(8.32)_=21.09, P<0.001, RM ANOVA, P<0.001 for each value, Tukey’s post hoc test as compared to the baseline). The percentage of fragmented mitochondria in the imaging field has recovered to 12.6±12.6% and 17.2±9.6% before the second and the third SD, respectively (P=1.0 for both values, Turkey post hoc test as compared to the baseline). Mitochondrial fragmentation during SDs was paralleled by SD-induced dendritic beading. As expected, the fraction of dendritic beading in the imaging field has picked at ∼100% during SDs (χ^2^ =31.3, P<0.001, Friedman RM ANOVA on Ranks) reaching 100% (IQR: 100-100%) during the first SD, 100% (IQR: 96.9-100%) during the second and 98.7% (IQR: 97.4-100%) during the third SD as compared to the baseline value of 2.4% (IQR: 0-9.4%) before the first SD (P<0.05, SNK post hoc tests). Dendrites completely recovered between SDs with a fraction of dendritic beading in the imaging field, returning to 1% (IQR 1-8%) before the second SD and 3.2% (IQR: 2.9-5%) before the third SD (P>0.05, SNK post hos test).

**Figure 2.**
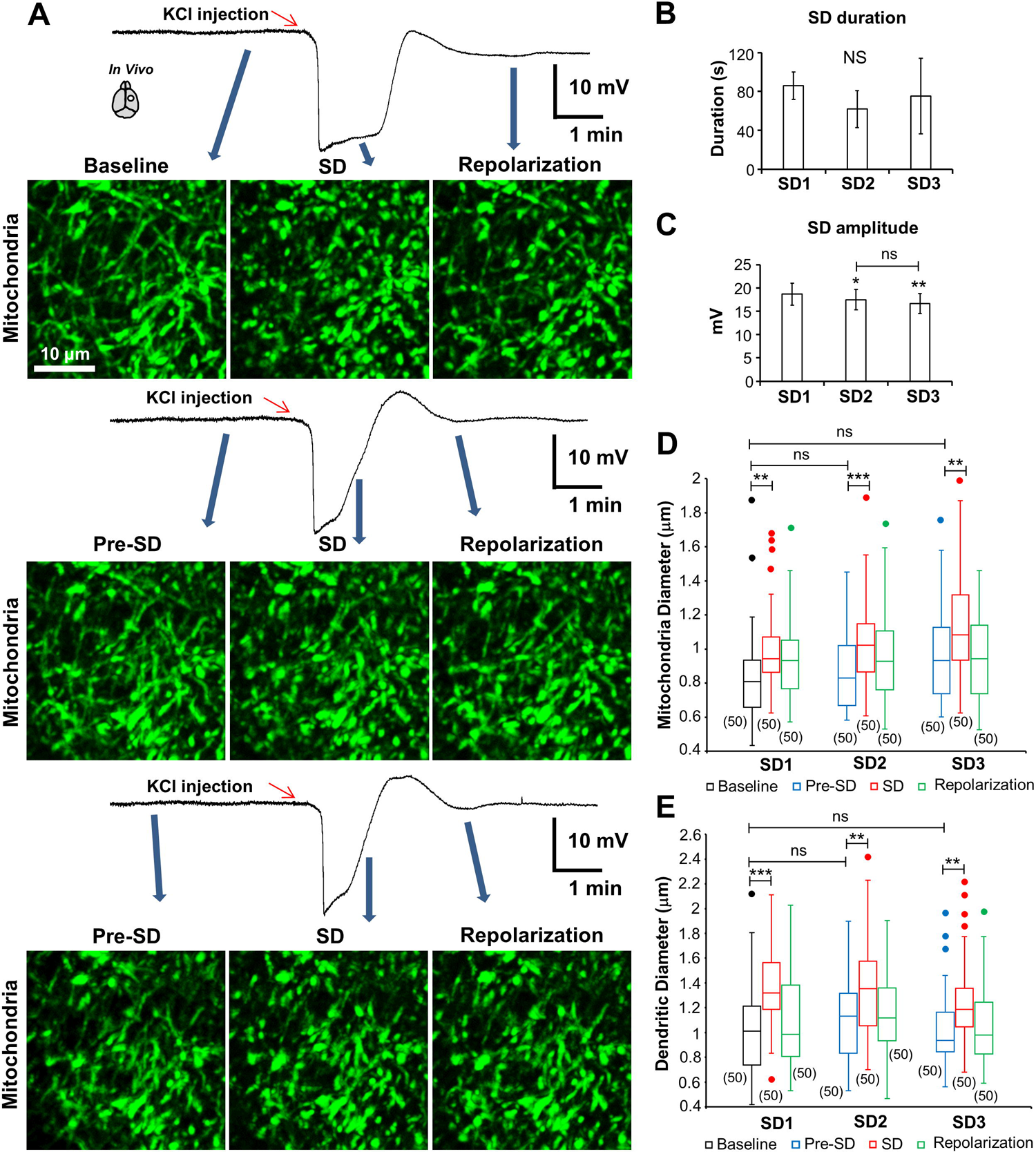
Several rounds of SDs do not cause irreversible mitochondrial fragmentation in the normoxic healthy neocortex. **A**, 2PLSM image sequence of the roGFP-positive mitochondria showing rapid fragmentation and recovery of a mitochondrial structure during the passage of three normoxic SDs induced with focal KCl microinjection away from the imaging field. Arrows correspond to various time points on SD traces recorded with a glass microelectrode placed next to the imaged mitochondria. 2PLSM image sequence of the same mitochondria exhibits a similar fragmentation pattern with a complete recovery before the second and third SD elicited at ∼33 min time intervals after the first SD. **B, C,** Quantitative features of 15 SDs from five mice. SDs were similar in duration (time between the points corresponding to half the DC amplitude and the same potential during recovery) (F_(2,8)_=1.59, P=0.26, RM ANOVA). SD amplitudes (potential difference between the start of SD and the lowest point of the DC deflection) have diminished by 6.2±4.4% for the second and by 10.8±4.5% for the third SD (F_(2,8)_=17.38, P<0.005, RM ANOVA with Tukey’s post hoc test. **P<0.005, *P<0.05 as compared to the first SD). **D,** Summary of changes in the diameter of 150 randomly selected mitochondria during three rounds of subsequent normoxic SDs in 5 mice. Quantification shows that mitochondria undergo similar rounds of transient swelling and recovery during the passage of three SDs evoked about every 23 min (χ2_(8)_=76.12, P<0.0001, Friedman RM ANOVA on Ranks with Dunn’s post hoc test. ***P<0.0001, **P<0.01, as compared with mitochondria diameter before each SD). Circles depict outliers. Numbers in parentheses indicate the number of measured mitochondria. **E,** Quantification of a diameter of 150 randomly selected dendrites during the passage of three rounds of subsequent normoxic SDs show transient dendritic beading/swelling (χ2_(8)_=108.83, P<0.0001, Friedman RM ANOVA on Ranks with Dunn’s post hoc test. ***P<0.0001, **P<0.001, as compared with dendritic diameter before each SD). Circles depict outliers. Numbers in parentheses indicate the number of dendrites included in the measurements.

Next, we corroborated these findings by measuring the diameter of randomly selected mitochondria and dendrites during the passage of subsequent SDs. Mitochondria significantly swelled during each SD (**Fig. 2D**; P<0.0001, Friedman RM ANOVA on Ranks), with diameter increasing by 13.3% (IQR: 3.7-28.4%) by 31.8% (IQR: 2.9-53.6%) and by 30.1% (IQR: 12.2-58.1%) relative to the baseline for the first, second and third SD respectively (P<0.01 for each value, Dunn’s post hoc test, as compared to baseline). Mitochondria diameter recovered to −0.1% (IQR: −19.7 – 22.5%) of baseline before the second SD and to 12.0% (IQR: −11.4 – 35.2%) of baseline before the third SD (**Fig. 2D**, P>0.05, Dunn’s post hoc test). Dendritic diameter increased 28.6% (IQR: 15.8 – 52.3%), 31.8% (IQR: 2.9 – 53.6%), and 15.8% (IQR: 2.0 – 32.1%) during three subsequent SDs (**Fig. 2E**, P<0.001, Friedman RM ANOVA on Ranks, P<0.001 for each measure, Dunn’s post hoc test as compared to baseline). The dendritic diameter has recovered to 10.2% (IQR: −18.7 – 28.2% and −8.9% (IQR: −17.7 – 13.4%) of baseline before each SD (**Fig. 2E**, P>1.0, Dunn’s post hoc test).

These data indicate that parallel to the innocuous normoxic SD-induced dendritic beading and recovery, mitochondria swell and recover during the passage of several rounds of SD without accumulating mitochondrial fragmentation.

### Dendritic mitochondria structure was affected by normoxic SD, as corroborated with ssEM

To confirm that mitochondrial fragmentation and swelling during SD were associated with substantial ultrastructural changes of mitochondria and not a redistribution of GFP, we conducted ssEM analyses in mice perfusion-fixed for EM immediately after confirmation of normoxic SD-induced mitochondrial fragmentation (“*SD*” group) (**Fig. 3A-C**). Since quantitative analyses of 2PLSM imaging data revealed that mitochondria were still recovering at 98.1 ± 14.4 s of SD onset (**Fig. 1D, E**), we attempted to perfusion-fix mice with aldehydes after detecting SD-induced mitochondria fragmentation within less than 2 min of SD onset (**Fig. 3B-C**). These experiments were exceptionally technically challenging, and only two mice out of 23 were used for quantitative ssEM analyses (<10% success rate). The rest of the mice were discarded, mainly because of the inability to perfusion-fix an animal within 2 min of SD onset. The labor-intensive nature of the ssEM analyses imposes excellent tissue preservation as a prerequisite for all EM samples; therefore, some tissue was also discarded based on this criteria. As a control, two “*Sham*” mice without SD were perfusion-fixed after baseline 2PLSM imaging over the somatosensory cortex and verification of the intact dendritic structure. To guide EM analyses to the area of 2PLSM imaging where mitochondria fragmentation was detected, we used the previously described method^41, 42^ to create fiducial ‘branding’ marks in fixed tissue using a Ti:sapphire laser beam (**Fig. 3D, E**). These laser marks were visible in the resin-embedded section and helped to identify the region of interest for serial sectioning within the area of 2PLSM intravital imaging (**Fig. 3F**). To reduce bias, all 292 dendritic mitochondria from 150 dendritic segments contained in two serial sections from two mice of the *SD* group were reconstructed in 3D and compared to 142 3D reconstructed mitochondria from 73 dendritic segments in three serial sections from *Sham* mice. As previously reported for dendritic mitochondria^22, 36, 37, 47^, *Sham* mice had mostly long filamentous mitochondria (**Fig. 3G**). In contrast, mitochondria from the *SD* group appeared shorter and swollen (**Fig. 3H**).

**Figure 3.**
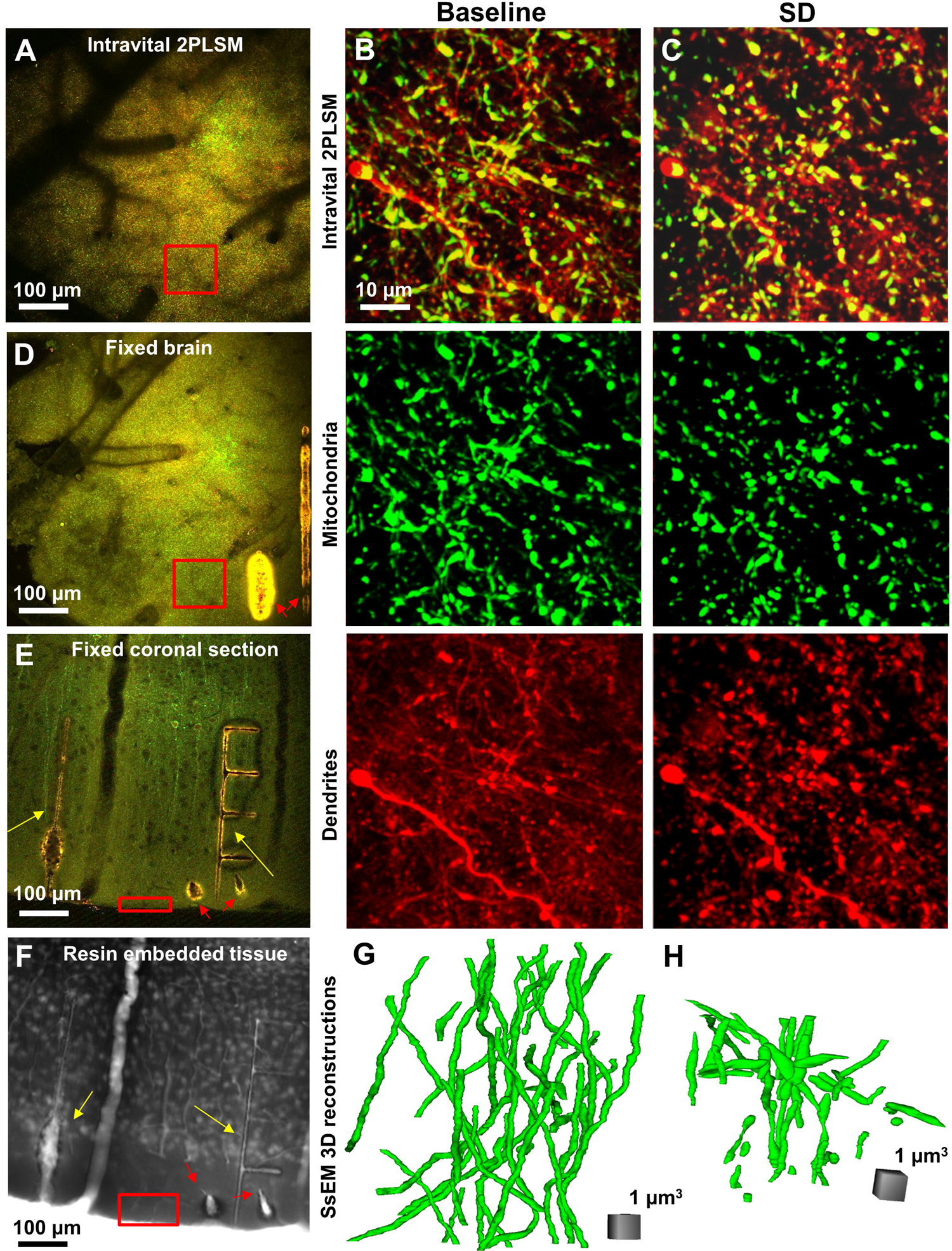
*In vivo* 2-photon imaging followed by quantitative ssEM of mitochondrial fragmentation during normoxic SD. **A,** *In vivo* low-magnification large-scale 2PLSM image of mouse somatosensory cortex expressing mito-roGFP-T2A-tdTomato construct (yellow). Dark structures are unlabeled vascular channels. The red box marks the area of high-magnification 2PLSM imaging. **B-C,** Pair of dual-color zoom-in-images of mitochondria (green) and dendrites (red) acquired within the red-boxed area in (A) before (B) and during SD (C). Split green and red channels at the bottom demonstrate mitochondrial fragmentation and dendritic beading coinciding with SD. After imaging, a mouse was perfusion-fixed through the heart within 2 min of SD onset. **D,** Low-magnification image of perfusion-fixed brain embedded in an agar block showing an area of intravital 2PLSM imaging (red box) identified using the blood vasculature and the dendritic pattern as a guide. Fiducial laser brands were created on the cortical surface (red arrows). **E,** Coronal section (100 μm thick) with visible original laser marks as in (D) (red arrows) and additional laser brands (yellow arrows) placed with the *in vivo* imaging area (red box) halfway between them. **F,** Laser branded tissue in resin processed for EM with clearly visible original laser marks (red arrows) and additional marks (yellow arrows) used to identify the location of *in vivo* 2PLSM imaging (red box). **G,** Example of 43 dendritic mitochondria reconstructed in 3D from ssEM in *Sham* mouse. **H,** Example of 43 mitochondria reconstructed in *SD* mouse from the same imaging field shown in (C).

### Quantification through ssEM

The length and maximum width of mitochondria were measured in 3D reconstructions. Mitochondrial segments from the *SD* group were significantly shorter (**Fig. 4A**; P<0.0001, K-S test), suggesting fragmentation. In *Sham* mice, the mitochondrial length was 5.16 (IQR: 2.0 – 8.98) μm compared to 2.62 (IQR: 1.48 – 4.52) μm in the *SD* group. However, these numbers underestimate the mitochondrial length as only 35.9% of mitochondria in *Sham* and 50% in the *SD* groups were entirely contained within the series, with the rest being cut at one or both ends of the series. It should be noticed that the proportion of “cut” mitochondria was not significantly different between conditions (χ^2^ =1.94, P=0.16) or between series (χ^2^ =4.97, P=0.29), indicating that the number of “cut’ mitochondria was not a confounding factor that influenced the difference in mitochondrial length between *Sham* and *SD* mice. Likewise, the maximum dendritic mitochondria diameter in the *SD* group was significantly larger by about 29%, indicating swelling and structural alterations (**Fig. 4B**; P<0.0001, K-S test).

**Figure 4.**
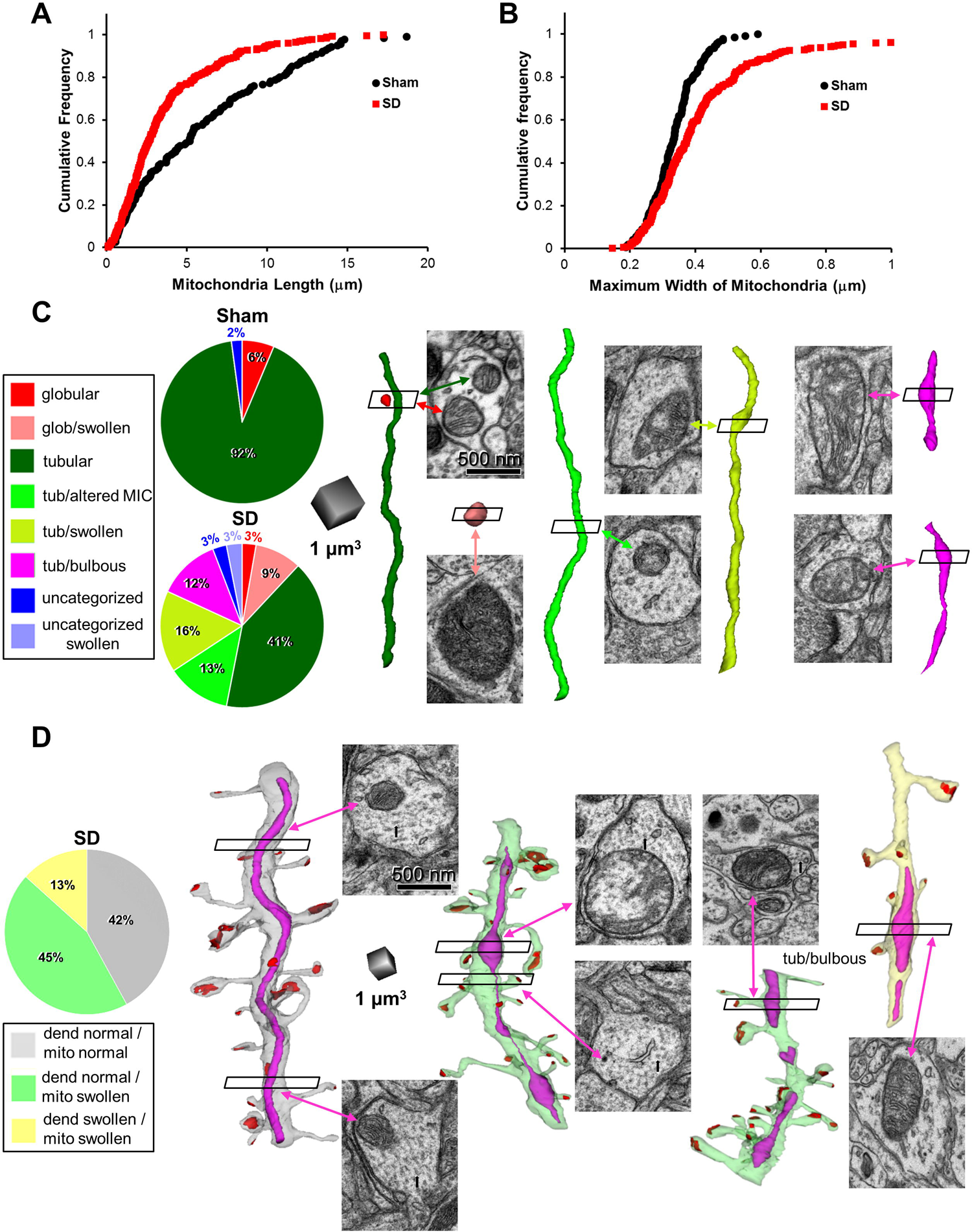
Alterations in dendritic mitochondria structure during normoxic SD. **A**, Cumulative frequency of length of each mitochondrial organelle in *Sham* and *SD* EM datasets. Mitochondria were significantly shorter in *SD* than in *Sham* mice (P<0.0001, K-S test). **B,** The maximum width of mitochondria was significantly larger in *SD* than in *Sham* groups (P<0.0001, K–S test). **C,** Morphological characterization of mitochondrial organelles in *Sham* and *SD* mice. Pie charts indicate the proportions of mitochondria in each category. Representative 3D reconstructions of mitochondrial organelles are color-coded according to the classification presented in pie charts. Sample EMs are positioned next to 3D reconstructions to illustrate mitochondrial ultrastructure in individual sections in the series. Arrows point to mitochondria. Globular (red) and tubular (green) mitochondria with normal appearance had uniformly gray matrices and cristae with evenly thin widths. Mitochondria with altered MIC (lime) had nonhomogeneous matrices and more variable and wider cristae. Swollen globular (pink), swollen tubular (olive), and tubular bulbous (magenta) mitochondria were wider with very nonuniform matrices and some cristae damage. **D,** Recovery of dendritic and mitochondrial morphology after normoxic SD. Pie chart colors match the colors of the reconstructed dendritic segments with excitatory synapses (red) and mitochondria (fuchsia). Representative EMs along dendrites showing cross-sectioned dendrite and mitochondria (arrow). Not swollen, normally appearing dendrites (gray and lime) have intact cytoplasm with arrays of microtubules (black arrows in the micrographs) and intact (gray dendrite) or swollen mitochondria (lime dendrite). Swollen dendrite (yellow) has a watery cytoplasm without microtubules and largely swollen mitochondria.

Next, we classified mitochondria into globules and tubules, selecting the aspect ratio of 3 (length/maximum diameter) as an arbitrary threshold between these categories^22, 47^. As previously reported for dendrites of pyramidal neurons^22, 36, 37^, most reconstructed mitochondria in *Sham* mice were elongated tubular organelles (91.5% of the total), and globular mitochondria were infrequent (6.3% of the total) (**Fig. 4C**). The remaining 2.1% of mitochondria were not classified because they were cut off too short at the edges of the serial sections. The proportion of globular mitochondria did not change significantly in the *SD* group (12% of the total) (χ^2^ =2.24, P=0.13). However, 77% of these globular mitochondria were swollen (globular/swollen) (**Fig. 4C**). The tubular structure with an unswollen 3D appearance was retained by only 53.8% of mitochondria in the *SD* group (χ^2^ =11.1, P<0.001, as compared to *Sham* mice). Yet, as assessed from individual EM sections in the series, 23.6% of these mitochondria (or 13.7% of the total) still showed some signs of swelling as judged by alterations in the mitochondrial inner compartment (MIC) cristae or matrix morphology in one or several places along the length of the organelle (**Fig. 4C**; tubular/altered MIC). The rest of the elongated mitochondria in the *SD* data set were swollen tubules (tubular/swollen; 16.1% of the total) or swollen tubules with enlarged protruding segments (tubular/bulbous; 12.3% of the total) (**Fig. 4C**), appearing, in the most extreme case, as a “mitochondria-on-a-string” (MOAS)^48^ (**Fig. 4D**). The remainder of mitochondria were uncategorized swollen and unswollen organelles because their were cut too short. Overall, 53.4% of reconstructed mitochondria in the *SD* group were swollen, or MIC had signs of swelling.

Next, we used the ssEM data set to gauge recovery from dendritic and mitochondrial swelling after normoxic SD. At the time of fixation, i.e., within less than 2 min of the SD onset, only 13.3% of dendrites were swollen with a watery cytoplasm and microtubule loss (**Fig. 4D**). None of these swollen dendrites contained structurally intact mitochondria and all mitochondrial organelles were swollen. The remaining dendrites appeared not swollen with intact arrays of microtubules, but many contained distended vacuoles indicative of recovery from dendritic swelling. Either swollen mitochondria or mitochondria with altered MIC were present in 44.7% of structurally unimpaired dendrites, while morphologically unaltered mitochondria were found in 42% of structurally intact dendrites. These ssEM data confirm findings from intravital imaging that mitochondrial recovery from normoxic SD-induced swelling takes longer than dendritic recovery.

### Tissue hypoxia is not a mechanism of dendritic mitochondria fragmentation during normoxic SD

In the normal cortex, SD is accompanied by a short period of tissue hypoxia manifested by a sharp drop in brain tissue partial pressure of oxygen (tpO_2_) to anoxic levels^24, 49–51^. Therefore, we tested if brain tissue hypoxia triggers mitochondrial fragmentation. To induce brain hypoxia, we decreased O_2_ concentration in the inspired air of mechanically ventilated mice. It has been reported that spontaneous SDs occur if the O_2_ concentration in the inspired air is lowered from 21% (normal air) to 8%^24^.

Therefore, to avoid spontaneous SDs, we reduced O_2_ in the inspired air of six mice to ∼10 % (10.2% (IQR: 10.1-10.4%)). As a result, no spontaneous SDs occurred in our experiments. At the same time, brain tpO_2_, measured with Clark microelectrode in four mice, dropped rapidly, within 17 s (IRQ: 15.5-22.3), from 26.5±13.1 mmHg at baseline to 3.7±4.2 mmHg (p<0.05, paired t-test), indicating tissue hypoxia. Notably, the same level of severe tissue hypoxia that lasted for ∼2 min was reported in mice during normoxic SD^24^. Nevertheless, mitochondria and dendrites steadfastly maintained their structure in the next 25 min (IQR: 10.25-31.5) of hypoxia (**Fig. 5A, B**) until terminal SD was triggered by a sudden cardiac arrest (CA). This terminal SD rapidly fragmented mitochondria and beaded dendrites (**Fig. 5C**). Indeed, quantitative analysis confirmed that the fraction of fragmented mitochondria in the imaging field significantly increased to 96.8±4.1% at CA coinciding with a terminal wave of SD (F_(2,10)_=1351.7, p<0.001, RM ANOVA with Tukey’s post hoc test). There were no changes in the percentage of fragmented mitochondria during hypoxia (6.5±0.9%) as compared to the baseline (8.6±4.9%) (P=0.56, Tukey’s post hoc test). Likewise, terminal SD beaded dendrites increasing the amount of beading to 100% (IRQ: 99.4-100%) (χ^2^ =8.33, P<0.05, Friedman RM ANOVA on Ranks with Dunn’s post hoc test), while the percentage of beaded dendrites was unchanged from the baseline value of 4.4% (IRQ: 2.5-6.1%) to 7.7% (IRQ: 6.9-9.5) during hypoxia (P=0.44, Dunn’s post hoc test).

**Figure 5.**
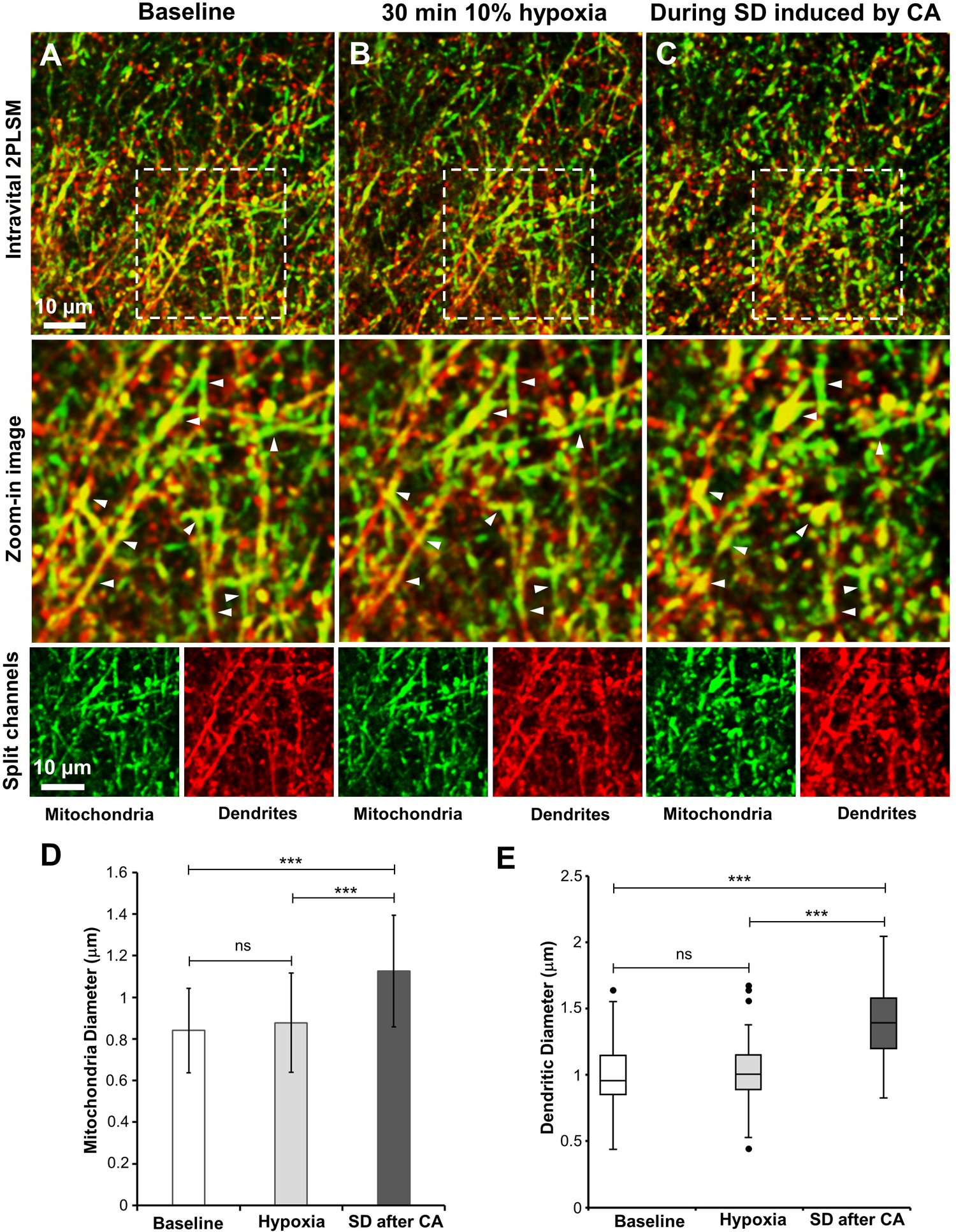
Mitochondria and dendrites remain intact during hypoxia until terminal SD after sudden cardiac arrest (CA). **A-C**, A large-scale dual-color 2PLSM MIP image sequence of dendrites and mitochondria in the top row with corresponding zoom-in images of areas in white boxes at the middle and images of mitochondria (green) and dendrites (red) as separate channels at the bottom demonstrate a lack of structural changes during 30 min of hypoxia (B) followed by rapid mitochondrial fragmentation and dendritic beading caused by terminal SD (C). Arrowheads point to the same spots before and during hypoxia and after terminal SD. **D,** Quantification of changes in the diameter of mitochondria during hypoxia and terminal SD showing significant swelling of mitochondria only at SD onset (F_(2, 118)_=76.51, P<0.001, RM ANOVA with Tukey’s post hoc test. ***P<0.001). **E,** Quantification of changes in the diameter of dendrites during hypoxia and terminal SD showing significant dendritic beading/swelling caused by SD (χ^2^ =58.43, P<0.001, Friedman RM ANOVA on Ranks with Dunn’s post hoc test. ***P<0.001).

Measurements of the diameter of the same 60 mitochondria in the six mice revealed mitochondria swelling caused by terminal SD during CA (**Fig. 5D,** P<0.001, RM ANOVA with Tukey’s post hoc test). Indeed, mitochondrial diameter increased by ∼34% during terminal SD (P<0.001 Tukey’s post hoc test) without any change during hypoxia alone (P=0.3, Tukey’s post hoc test). The difference in the diameter of the same 60 dendrites, measured in the six mice, also revealed significant dendritic beading/swelling by ∼39% during terminal SD (**Fig. 5E**, P<0.001, Friedman RM ANOVA on Ranks with Dunn’s post hoc test). However, the dendritic diameter was unaffected by hypoxia (P=0.6, Dunn’s post hoc test). These results have shown that dendrites and their mitochondria remained structurally intact during ∼25 min of hypoxia but were rapidly affected by terminal SD during CA.

## Discussion

This study is the first to characterize the neuronal mitochondrial structural dynamics *in vivo* during normoxic SD. In the non-injured neocortex, the SD wave, which is thought to underlie migraine aura, results in transient dendritic mitochondria swelling and fragmentation alongside dendritic beading. Mitochondrial structure recovered, but recovery took longer than recovery of dendritic structure. Several rounds of normoxic SD did not result in the accumulation of mitochondrial fragmentation or terminal dendritic swelling, which agrees with the previous report of reversible dendritic beading during normoxic SD^27, 29^. Surprisingly, the dendritic mitochondrial structure *in vivo* was resilient to energy deficits during severe hypoxia until terminal SD induced by cardiac arrest.

Neuronal mitochondria structure is very sensitive to a full spectrum of cortical injuries, as recently revealed by *in vivo* real-time 2PLSM imaging of mitochondrial dynamics^37^. Not surprisingly, a single SD elicited by severe energy deficits during transient global ischemia coincided with neuronal mitochondria fragmentation^22^.

Likewise, on the other end of the SD continuum, normoxic SD in healthy, well-nourished cortex also triggered rapid mitochondrial fragmentation. Mitochondrial fragmentation was not strictly dependent on the energy status of the cortical tissue as mitochondrial organelles morphology was not affected by severe tissue hypoxia for many minutes until the arrival of the terminal SD after cardiac arrest. These findings support the idea that SD is the important triggering mechanism of rapid neuronal mitochondrial fragmentation along the full spectrum of noxious conditions evoking SD.

Mitochondrial structure underlies its function^52^: thin and long dendritic mitochondria are healthy^22, 36, 37^, swollen and fragmented mitochondria are likely dysfunctional^53^. Under non-deleterious conditions, mitochondria are very dynamic organelles that undergo fusion and fission^46, 54, 55^. Such dynamics is necessary to retain a healthy mitochondrial population through fusion, which is thought to maintain mitochondrial bioenergetics, and fission, which partitions injured mitochondria from the healthy mitochondrial network for degradation by mitophagy^53, 56, 57^. SD is metabolically taxing^10^, and even in the healthy neocortex, the energy demand of Na^+^/K^+^-ATPase increases so markedly^58^ that the tissue ATP concentration falls to about 50% of the baseline level^59, 60^. It is imaginable that normoxic SD-induced mitochondrial fission could boost ATP production because, under certain experimental conditions, fragmented dendritic mitochondria could produce more ATP on acute demand^61^. However, under noxious conditions, fragmentation of the mitochondrial network into small spherical structures is considered a hallmark of mitochondrial injury^46, 62^. Importantly, excessive mitochondrial fragmentation also causes increased production of reactive oxygen species and release of pro-apoptotic factors leading to cell death^53, 54, 63–66^. Albeit several successive normoxic SDs did not result in irreversible mitochondrial fragmentation, it is plausible that in the metabolically compromised penumbra, SD-induced mitochondrial fragmentation is not readily reversible, with each subsequent SD exacerbating persistent fragmentation. Sustained excessive mitochondrial fragmentation is likely incompatible with cell survival and might signify the “commitment” point for neurons to die^2, 67^. It is conceivable that the imbalance between mitochondrial fragmentation and fusion along the SD continuum is one of the key mechanisms underlying the SD-imposed transition to cell death and warrants future investigation.

Low oxygen levels during hypoxia should inhibit mitochondrial oxidative phosphorylation^68^, which slows Na^+^/K^+^-ATPase and ignites SD^10^. Still, we could avoid spontaneous SDs by keeping oxygen level in the inspired air at ∼10% with parallel tpO_2_ levels in the range of severe tissue hypoxia closely corresponding to the tpO_2_ levels reported during normoxic SD^24^. During severe energy deprivation, neurons eventually swell even without SD because of the cation influx driven by Gibbs–Donnan forces^23, 69^ and inadequate cation outflux caused by slowing down ATP-dependent sodium and calcium pumps. Nevertheless, dendrites maintain their structure, indicating sufficient ATP produced by glycolytic and residual oxidative metabolism^70^. Inhibition of mitochondrial oxidative metabolism with chemical treatment leads to mitochondrial fragmentation^71, 72^. Mitochondria maintained their structure in our hypoxic conditions, possibly reflecting an adaptive shift in the state of the mitochondrial network toward fusion to keep ATP generation while inhibiting fragmentation^73, 74^. Regardless of the mechanism, our findings indicate that energy deprivation during normoxic SD is not a mechanism of mitochondrial fragmentation.

Mitochondrial fragmentation and swelling during normoxic SD first observed with real-time 2PLSM were corroborated with quantitative ultrastructural analyses. Although we were unable to reconstruct specifically the same mitochondria and dendrites observed with 2PLSM, we had precisely collected ssEM samples from the 2PLSM imaging field guided by laser branding^41, 42^. One limitation is that ssEM data were acquired only from two animals. The low number of mice was due to a very challenging requirement to perfusion fix mice in less than 2 minutes after SD onset, i.e., during repolarization or immediately after, when a substantial fraction of mitochondria was still swollen and fragmented, and some dendrites were still recovering from beading.

Despite this apparent technical difficulty, ssEM data provided a snapshot of recovery from normoxic SD-induced brain edema, signified by some swollen dendritic profiles with watery cytoplasm, loss of microtubules, and swollen organelles. Previously, only one EM study examined cellular swelling during normoxic SD using a rapid freeze-substitution technique^75^. Despite the neuropil disruption by ice crystals, analyses of a few superficial micrometers of the cortex were possible. This study revealed transient swelling of dendritic profiles elicited by normoxic SD. The mitochondrial morphology was not analyzed due to technical limitations. Here, using aldehydes fixation, which affords excellent tissue preservation, we have shown that 53% of mitochondria and 13% of dendrites were swollen or had signs of swelling at the time of fixation. Also, 45% of normally-appearing dendrites that apparently recovered from swelling still had swollen mitochondria. These EM data verified 2PLSM analyses of dendrites and their mitochondria, which showed that although mitochondrial fragmentation temporally coincided with dendritic beading, mitochondrial recovery was slower than recovery of dendritic morphology. These data also agree with our previous *in vivo* 2PLSM demonstration of heterogeneous sensitivity to the injury between dendrites and their mitochondrial organelle^37^.

In highly artificial systems such as cultured neurons, water influx during hypo-osmotic stress was sufficient for mitochondrial collapse into rounded swollen structures^76^. Contrary to cultured neurons, intact neurons are osmo-resistant^77^ due to the lack of aquaporins in their plasma membrane^78^ but swell during SD. Dendritic swelling elicited by SD results from water accumulation^22, 27, 77, 79^. It is possible that SD-induced mitochondrial fragmentation is caused by water influx. EM datasets showing the absence of swollen dendrites with intact mitochondria might indirectly support this idea. Conversely, despite astrocytes swell during SD^20, 69^, astroglial mitochondria maintaine their structure during ischemia-induced SD, which is in striking difference to the neighboring dendritic mitochondria^47^. Whether SD-induced water influx into intact neurons is sufficient and necessary for neuronal mitochondria fragmentation requires further investigation.

Several other mutually not excluding mechanisms can be proposed to link SD across the full spectrum of SD waves and noxious conditions to the collapse of the neuronal mitochondrial network. SD-induced mitochondrial fragmentation may result from mPTP opening^80–82^ or disruption of mitochondria dynamics^53^ or both. During SD, dendritic Ca^2+^ rises from a resting level of ∼80 nM^83, 84^ to ∼20 μM^85^. Mitochondria rapidly take up Ca^2+^ through mitochondrial uniporter^86^ to prevent Ca^2+^ overload into the cytosol^87, 88^. During SD, excessive Ca^2+^ absorbed by mitochondria might result in mitochondria Ca^2+^ overload, leading to mitochondrial permeability transition pore (mPTP) induction^81, 89–91^. Ca^2+^-dependent mitochondria transformation from elongated to swollen and rounded structures has been linked to mPTP induction in neuronal cultures^92^ and in vivo^35^. Indeed, the blockade of mPTP prevented^92, 93^, or blunted these morphological changes^35^. Future experiments are warranted to address mechanisms of SD-induced mitochondrial fragmentation.

In conclusion, we uncovered that even a single normoxic SD, which is thought to underlie migraine aura, fragments the neuronal mitochondrial network, likely affecting mitochondrial organelle bioenergetics. Consequently, SD is a common mechanism eliciting mitochondrial fragmentation. Mitochondrial fusion occurs rapidly during repolarization from normoxic SD. Like dendritic beading that reflects the onset of SD-induced neuronal cytotoxic edema along the entire SD continuum, mitochondria fragmentation is similarly elicited along the SD spectrum. Since mitochondrial structural remodeling affects neural pathological outcomes, it is feasible that therapeutic approaches targeting mitochondrial dynamics in neurological conditions triggering SD will be beneficial to improve cell survival and limit SD-induced injury.

## Author contributions

SAK designed the study; JS performed experiments and analyzed 2PLSM data; IVF conducted EM analyses; SAK participated in 2PLSM and EM data analyses and wrote the manuscript. All authors edited and approved the manuscript before submission.

## Acknowledgments

We thank Libby Perry and Brendan Marshall (Electron Microscopy Core at the Medical College of Georgia) for their assistance with electron microscopy.

## Data availability statement

The data supporting this study’s findings are available from the corresponding author upon reasonable request.

